# Estimation of the cancer risk induced by rejuvenation therapy with young blood and treatment recommendations

**DOI:** 10.1101/298836

**Authors:** Michael Meyer-Hermann

## Abstract

In recent years the transfer of blood from young to old individuals was shown to bear the potential of rejuvenation of stem cell activity. While this process might increase life expectancy by prolonging functionality of organs, higher cell replication rates bear also the risk of cancer. The extent of this risk is not known.

While it is difficult to evaluate this cancer risk in experiments, this is possible with a mathematical model for tissue homeostasis by stem cell replication and associated cancer risk. The model suggests that young blood treatments can induce a substantial delay of organ failure with only minor increase in cancer risk. The benefit of rejuvenation therapy as well as the impact on cancer risk depend on the biological age at the time of treatment and on the overall cell turnover rate of the organs. Different organs have to be considered separately in the planning of the systemic treatment. In particular, the model predicts that the treatment schedules successfully applied in mice are not directly transferable to humans and guidelines for successful protocols are proposed. The model presented here may be used as a guidance for the development of treatment protocols.

**Additional information:** There is NO competing interests.

---

Biological organisms like humans accumulate DNA damage in replicating cells inducing many processes summarised under the general term of ageing. Without DNA damage, stem cell activity could be maintained on a level that guarantees better homeostasis of the organism. However, stem-cell function declines over time and is associated with ageing [1]. There is increasing evidence that the process of decline of stem-cell function can be reverted by transfer of blood from younger individuals in liver [2] and brain [3]. This was shown with heterochronic parabiosis mouse models [4, 5], in which young and old mice share their circulatory system.

It is becoming increasingly clear that it is not the cellular components of the young blood but soluble factors that mediate the positive effect on stem-cells in various organs including brain, liver, and muscles [6]. The determination of those systemic factors responsible for reactivation of stem-cells is ongoing [7]. More recent data suggest that it is not a property of young blood, but the dilution of inhibitory factors in old blood that would be responsible for the effect [8].

Irrespective of the actual mechanism that induced improved tissue homeostasis, reactivation of damaged and silenced stem cells might well induce a higher risk of developing age-related diseases like cancer [9]. While it is impossible to quantify the degree to which stem cell re-activation would increase the risk of cancer in experimental settings, this is possible based on theoretical methods and is the main objective of the present research. With a minimum of assumptions the presented analysis predicts the price of a stem cell rejuvenation therapy in terms of the impact of the therapy on cancer risk.

## Results

### A simple and generic model of organ homeostasis and cancer development

In order to ensure predictive power of the mathematical model it has to be constructed on the basis of rather generic assumptions. The model complexity is chosen such that the driving question, whether cancer risk is under control during rejuvenation therapy or not, can be addressed, avoiding any speculation on molecular mechanisms. The model (see Methods: *Model equations and analytical solution*) distinguishes stem (*S*) and tissue cells (*T*). It further distinguishes two reasons for death: organ failure and cancer. More specifically the model is based on the following set of assumptions:

- Tissue cells (*T*) have a natural life time (death rate *δ*) and are replenished by stem cell (*S*) division (proliferation rate *p*).
- The ratio of stem to tissue cells is 1:1000.
- Humans have a life time (Θ) of 80 years, mice of 2 years.
- Death is induced by organ failure, which happens when the total number of tissue cells hits a fraction (*f*) of 50%.
- Both, tissue and stem cells accumulate damage (mutation rate *γ*). The kind of damage is not distinguished and only the number of accumulated damage events is considered.
- Damaged stem cells pass their damage state to tissue progeny in the course of homeostatic division.
- The damage rate (*γ*) is constant.
- Cancer is induced at a critical number of mutations per cell (*c*).
- Cancer risk at the age of 80 is *ξ* = 30%.

The rate of point mutations in human adult stem cells was measured in the range of 40 mutations per year [10], where an approximately linear relationship of the number of mutations over age was found. This supports the assumed constant damage rate (*γ*). While the number of stem cell mutations correlates with cancer incidence [11], this absolute frequency of mutations is still difficult to be related to cancer risk because not all of these mutations happen in oncogenic regions of the genome. Furthermore, the immune system is removing pathogenic cells, which makes the absolute number of mutations associated with cancer unclear. In order to keep the model simple, I relied on the linear accumulation of mutations over age and fixed the cancer-inducing number of mutations to *c* = 10 per cell. This choice can be considered as a worst case scenario for the cancer risk induced by rejuvenation therapy because, on the one hand, cancer is expected to be induced by more than 10 accumulated mutations, and, on the other hand, with *c >* 10 the induced cancer risk is getting smaller in the model, as was revealed by a robustness analysis (see Methods: *Robustness of the statements*).

As the division (*p*) and the cell damage rate (*γ*) are determined by the life expectancy (Θ; see Methods: *Determination of the division rate*) and the cancer risk (*ξ*; see Methods: *Determination of the mutation rate*) at this age, respectively, this model has only a single free parameter: The cell death rate *δ*, which controls the overall turnover rate of tissue cells. The turnover rate is a tissue specific property and different values are considered in the following. Parameter sets for *low, intermediate*, and *high turnover* are defined in Table 1 and used as reference parameter sets.

**Table 1:** Reference model parameter sets. The rates *p*, *p*_max_, *δ*, and *γ* are given per year. Life expectancy Θ and half time *K* are in years. Note that *p* (Methods: *Determination of the division rate*), *p*_max_ (Methods: *Determination of the division rate of perfect homeostasis*), and *γ* (Methods: *Determination of the mutation rate*) are calculated from side conditions, thus, are not free parameters. The turnover rate is controlled by the cell death rate *δ*, where *low*, *intermediate*, and *high* turnover is used for simulations of humans. In runs with age-dependent *p*(*t*), the value for *p* is replaced by the three values *p*_max_, *K*, and *n* (see Eq. (15)), where *p*_*max*_ and *K* are calculated from side conditions (see Methods: *Age-dependent stem cell division rate*). Deviations from these sets are explicitly mentioned in the text. The robustness of the results against parameter variation is discussed in Methods: *Robustness of the statements*.

| turnover | $\sigma_0$ | $\tau_0$ | $\Theta$ | $f$ | $p$ | $\delta$ | $\gamma$ | $c$ | $\xi$ | $p_{\max}$ | $K$ | $n$ |
| --- | --- | --- | --- | --- | --- | --- | --- | --- | --- | --- | --- | --- |
| low | $10^5$ | $10^8$ | 80 | 0.5 | 3.41 | 0.014 | 0.0091 | 10 | 0.3 | 13.7 | 19.7 | 2 |
| intermediate | $10^5$ | $10^8$ | 80 | 0.5 | 27.2 | 0.055 | 0.0091 | 10 | 0.3 | 54.8 | 60.4 | 2 |
| high | $10^5$ | $10^8$ | 80 | 0.5 | 68.5 | 0.14 | 0.0091 | 10 | 0.3 | 137 | 72.4 | 2 |
| mouse | $10^5$ | $10^8$ | 2 | 0.5 | 105 | 0.5 | 0.32 | 10 | 0.1 | 500 | 0.4 | 2 |

Starting from a young individual without any damaged cell, the time course of cell damages over age is depicted in Figure 1. In this setting, the age of death is determined by the time, at which the total number of cells building up the virtual organ hits the fraction *f* = 50%. This time will be denoted as *age of organ failure* in the following. The age of cancer onset is determined by the time when the sum of all cells with *c* = 10 or more mutations passes the limit of 1 cell (purple lines in Figure 1). Assuming ergodicity, one can state that a number of cells with 10 or more mutations below 1 reflects the cancer risk at a particular age. In the following, the *age of cancer risk ξ* will be discussed, which denotes the age at which the number of cells with *c* = 10 or more mutations reaches *ξ* = 0.3.

**Figure 1:**
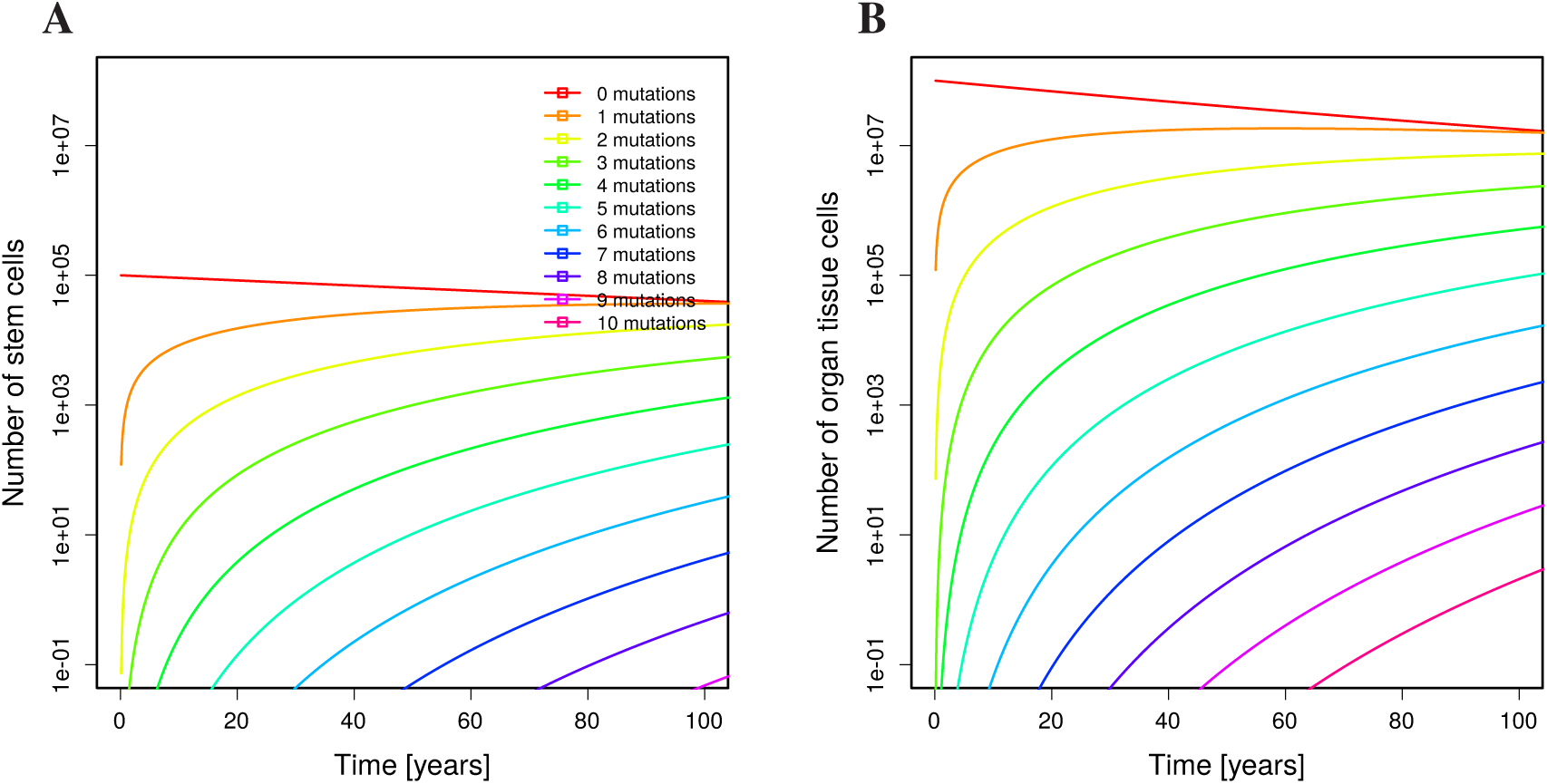
Damage dynamics in stem and tissue cells. Only undamaged stem (**A**) and tissue cells (**B**) exist in the beginning (initial conditions are *σ*_*i*_ = *τ*_*i*_ = 0 for *i >* 0 in Eq. (2)). Parameters of *low turnover* in Table 1 were used. The number of cells with different numbers of mutations over the life time is shown with different colours. The line with 10 mutations recollects all cells with *c* = 10 or more mutations.

### Rejuvenation therapy impacts on the age of organ failure

Blood transfer from young individuals is meant to reactivate stem cell division. In the model this is reflected by changing the stem cell division rate *p*. Figure 2 shows how the ages of organ failure Eq. (14) and cancer risk *ξ* Eq. (12) depend on the induced division rate, where the induced division rate is assumed constant over the life time. The untreated division rate determined in Eq. (6) (vertical dashed line), by construction, induces organ failure and a cancer risk of *ξ* = 0.3 at the age of Θ = 80 years. The division rate for perfect tissue homeostasis determined in Eq. (11) prohibits death by organ failure. Surprisingly, the result clearly shows that the age of organ failure is highly sensitive to the division rate (red line in Figure 2) while the age of cancer risk *ξ* is hardly changing (blue line in Figure 2). For a young blood therapy inducing stem cell replication to a degree that organ failure happens at the age of 400 (this is achieved by doubling the stem cell replication rate), the age at which the cancer risk reaches 30% is shifted from 80 to 78.

**Figure 2:**
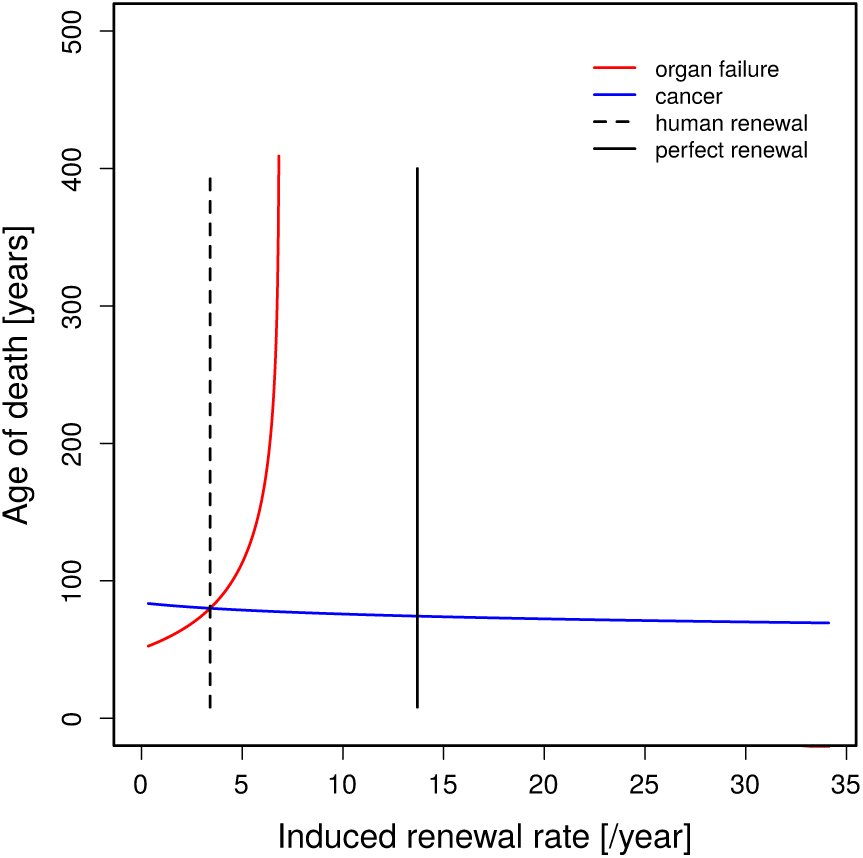
Ages of organ failure or cancer risk *ξ* in dependence on the cell renewal rate. The age of organ failure (red line, Eq. (14)) is highly sensitive to changes in the cell renewal rate (*p*) while the age of cancer risk *ξ* (blue line, Eq. (12)) remains comparably stable. Unmodified stem cell renewal *p*_normal_ (see Eq. (6)) and perfect cell renewal *p*_juvenile_ (see Eq. (11)) are marked by the vertical dashed and full lines, respectively. Between the red and the vertical black line, Eq. (14) has no solution, which means that organ failure never happens. Parameter set for low turnover in Table 1.

### Organ failure is delayed with limited cancer risk

A life long treatment with young blood is not a realistic scenario. Assuming a treatment limited to a duration of 10 years starting at different ages (see Methods: *Treatment at particular ages*), a similar picture in terms of sensitivity of the ages of organ failure and cancer risk emerges (full lines in Figure 3). The overall effect on the age of organ failures is smaller than for life long treatments (Figure 3A,B). With an induced division rate of perfect tissue homeostasis Eq. (11), organ failure happens at ages between 85 and 100 years, and the cancer risk of *ξ* = 30% is reached at ages around 79 ± 1 years. When the same treatment is repeated three times every 10 years with pausing intervals of 10 years (dotted lines, Figure 3), organ failure happens at ages between 110 and 150 years and the cancer risk of 30% is reached at the age of 78 ± 1 years. While the absolute numbers have to be interpreted carefully, the difference in the sensitivity of both read-outs is a stable finding.

**Figure 3:**
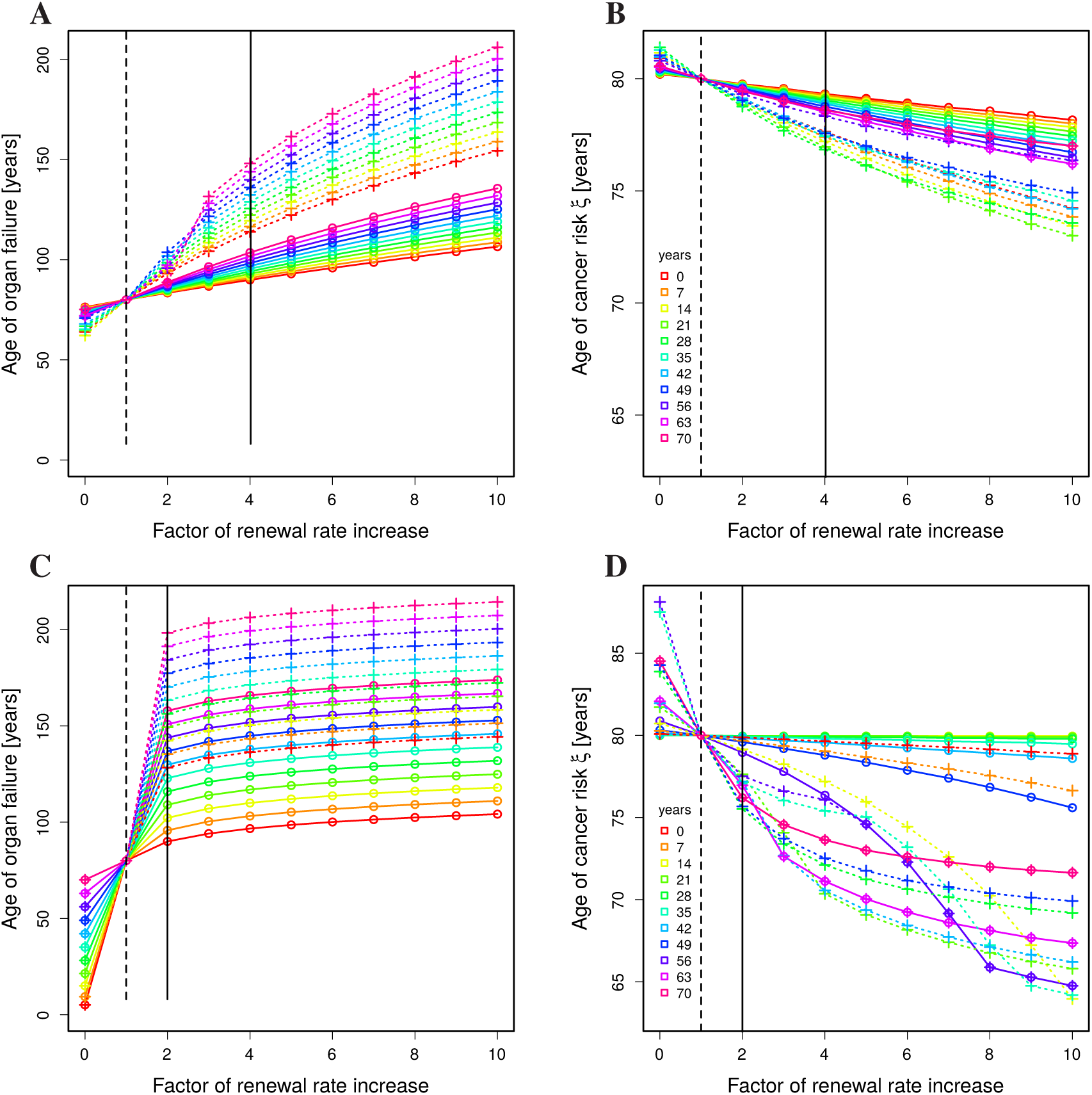
Organ failure and cancer risk with time-limited rejuvenation treatment. The analytical solution Eq. (2) together with 10 year treatments (Methods: *Treatment at particular ages*) is analysed for ages of organ failure (**A,C**) and cancer risk *ξ* (**B,D**) in dependence on the age of first treatment (colors) and with different induced renewal rate *p* (horizontal axes). A single treatment (full lines, circles) is compared to three treatments with interruptions of 10 years (dashed lines, crosses). Vertical lines denote the renewal rate of humans with a life expectancy of 80 years (dashed black line) and with perfect tissue homeostasis (full black line). Parameter set of low (**A,B**) and high (**C,D**) turnover in Table 1. Compare Supplementary Figure 1 for intermediate turnover rates.

Organ failure is delayed the more the later the treatment is started. Cancer risk is increased correspondingly. Surprisingly, when the first of the three treatment is started at an age of 42 or later, cancer risk is getting lower when the treatment is started later (see Figure 3A,B; blue and shorter wavelength curves). In this regime, rejuvenation treatment has a strong effect on the delay of organ failure with a comparably small increased risk of cancer. However, overall organ functionality is lower when treatment is started later. The treated individual faces a physiological state that is kept above but near of the organ failure threshold, which implies a lower life quality compared to treatments started at younger age.

### High-turnover organs are more sensitive to rejuvenation treatment but less stable

The investigation was built on organs with a low turnover rate (see Table 1, *low*). For high turnover organs (Figure 3C,D), the sensitivity of the age of organ failure to a single rejuvenation treatment is even higher. With an induced division rate of perfect tissue homeostasis Eq. (11), organ failure happens at ages between 90 and 160 years, while the age of cancer risk *ξ* is kept between 76 and 80 years. With three treatments of 10 years, the age of organ failure can be delayed to ages of 130 to 200 years, while the cancer risk is not further increased. Importantly, for high turnover organs, a single rejuvenation treatment can delay the age of organ failure to 120 years, while inducing no major change in the cancer risk (Figure 3C,D; full green and longer wavelength lines).

Failure of high turnover organs cannot be further delayed by higher than perfect homeostasis *p*_juvenile_ stem cell replication rates. However, cancer risk enters a non-linear regime at higher replication rates (Figure 3D). Organs with higher turnover rates react with higher or lower cancer risks than for low turnover organs depending on how and when the treatment is applied. There is hardly any increased cancer risk when the treatment is done at young age, irrespective of the target replication rate of the treatment. There is a transition to a strong impact on cancer risk at the age of 35 to 42. Cancer risk is highest for treatments started at the age of 56 and decreases again at higher ages (Figure 4). As the treatment is applied systemically, it will always also affect high turnover organs. The presented results illustrate that the timing and the dosage have to be planned quantitatively in order to avoid adverse effects.

**Figure 4:**
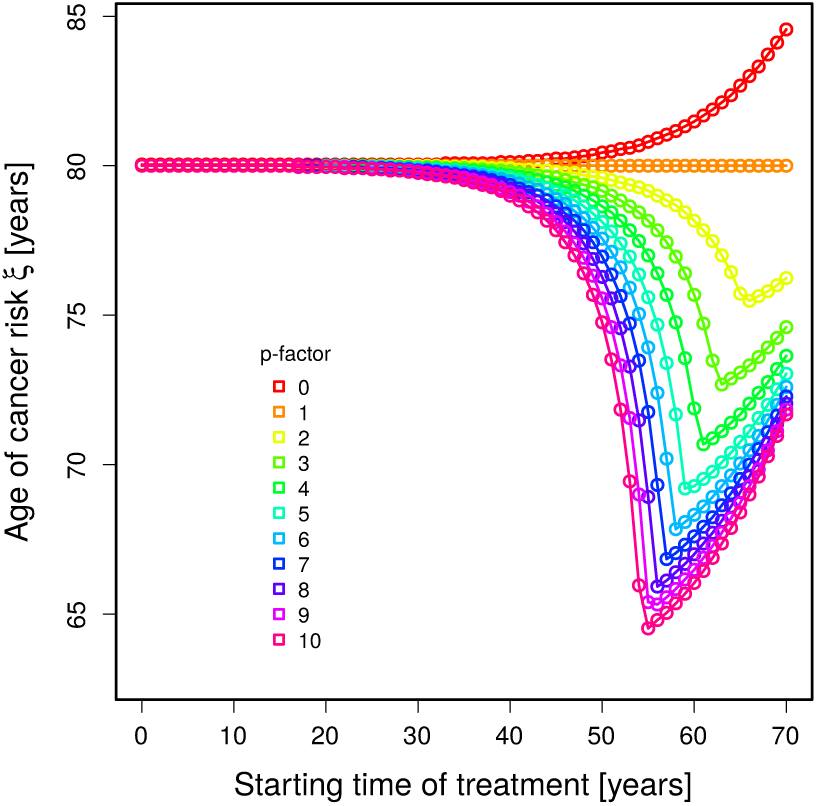
Optimal treatment age to minimise cancer risk in high turnover organs. Same data as in Figure 3C,D with a single 10 year treatment. The age of cancer risk *ξ* is shown in dependence on the time of treatment (horizontal axis) for different strength of treatment reflected by the p-factor (colors). Parameters of high turnover in Table 1.

### Limited cancer risk also in a model with age-dependent loss of stem-cell replication

A general tendency of rejuvenation treatment is that the later the treatment is started, the more organ failure is delayed and the more cancer risk is increased (Figure 3). However, this result might rely on the assumption that stem cell replication is constant over the life time but for the periods of treatment. While this assumption is a good approximation for skin tissue, in which stem cell replication is widely conserved at all ages [12], other organs exhibit a decrease of the stem cell replication rate with age [1]. For example, hematopoietic stem cells lose power in the course of ageing [13, 14]. The degree of this reduction is again organ specific. Assuming a Hill-function for the dependence of *p* on age (see Methods: *Age-dependent stem cell division rate*), the set of differential equations Eq. (1) cannot be solved analytically anymore and is solved numerically.

Induction of higher stem cell replication rates in low turnover organs exhibits a qualitatively similar result as with constant *p* (Figure 5A,B). The initial (at birth) replication rates required for organ failure at ages above 100 are higher because of the loss of replication with increasing age. An increase of the replication rate to an absolute level has a stronger effect, the later in life the treatment is done. The sensitivity to the treatment of the age of organ failure is still substantially higher than the sensitivity of the age of cancer risk *ξ*.

**Figure 5:**
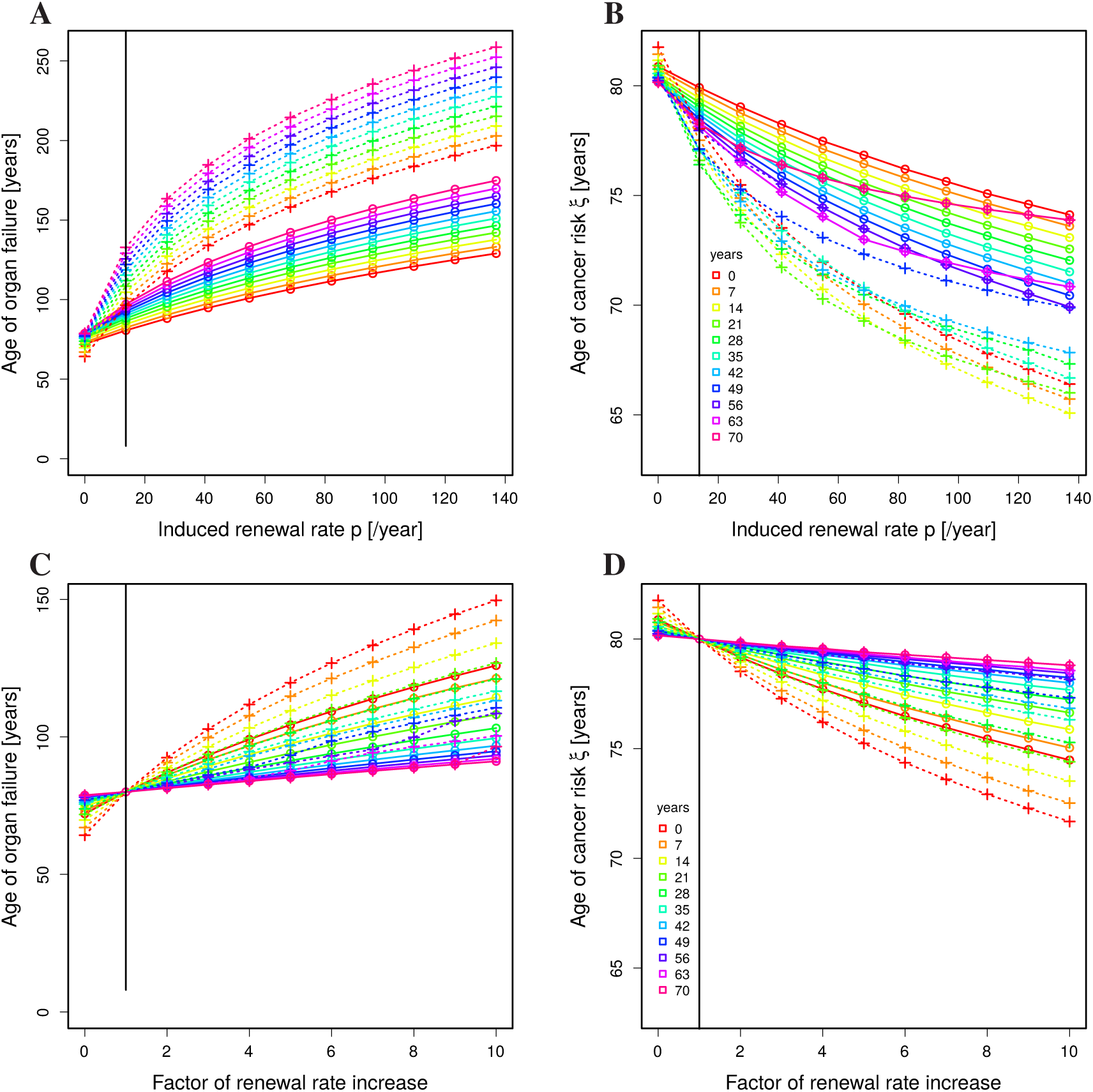
Absolute versus relative impact of rejuvenation treatment in a model with age-dependent loss of stem-cell replication. The numerical solution of Eq. (1) together with 10 year treatments (Methods: *Treatment at particular ages*) is analysed for ages of organ failure (**A,C**) and cancer risk *ξ* (**B,D**) in dependence on the age of first treatment (colors) and on the strength of treatment (horizontal axes). Absolute (**A,B**) or relative improvements (**C,D**) of the renewal rate are distinguished. A single treatment (full lines, circles) is compared to three treatments with interruptions of 10 years (dashed lines, crosses). Vertical lines denote the renewal rate *p*_*juvenile*_ inducing perfect tissue homeostasis (full black line). Parameter of low turnover in Table 1 with age-dependent stem-cell replication rate *p* in Eq. (15). For higher turnover rates compare Supplementary Figure 2 (absolute renewal rates) and Supplementary Figure 3 (relative renewal rates).

### The mechanism of rejuvenation therapy impacts on the hierarchy of best treatment age

Rejuvenation treatment with young blood at a particular age can either relatively improve the remaining stem cell replication activity or induce an absolute replication level. The mechanism of how young blood transfer improves stem cell replication is not known. Therefore, the aforementioned results, which were based on absolute renewal rates, are compared to results from a model with relative improvements of the renewal rate.

When treatment with young blood induces an improved replication relative to the level of stem cell replication at the time of treatment, the situation changes in organs with low turnover rates (Figure 5C,D). At first, the overall impact on the ages of organ failure and cancer risk *ξ* is smaller. Secondly, the advantageous age of treatment is different: With a factor of 10 higher stem cell replication in young age, a substantial retardation of organ failure can be induced. However, the later the treatment is done, the weaker the effect, thus, inverting the result found with either constant stem cell replication (Figure 3) or age-dependent stem cell replication treated to adopt absolute replication levels (Figure 5A,B). 10 times no replication is still a weak replication. At advanced age, a treatment with a relative improvement of stem cell replication has weak effects on both, the age of organ failure and the shift of cancer risk.

For organs with intermediate or high turnover (Supplementary Figure 3), the original sequence as found in (Figure 3 and Figure 5A,B) is restored, i.e. the later the treatment the more organ failure is retarded. As the sequence of whether treatment of young or old is more advantageous in terms of retardation of organ failure exhibits a switch dependent on the organ turnover level, an turnover rate may exist, at which the treatment effect becomes widely independent of the age of treatment. This is, indeed, the case (Supplementary Figure 4). This result suggests that, provided young blood therapy induces a relative increase of stem cell activity, the best age for treatment is different for different organs.

### Rejuvenation treatment optimised in mice cannot be transfered to men

In C57BL/6 mice with a mutation associated with the development of Alzheimer’s disease, parabiosis of young and old mice for 5 weeks could ameliorate cognitive deficits developed in the old mice [2, 3]. The mathematical model was challenged by testing whether an effect on age of organ failure would be found in this setting. The life expectancy of C57BL/6 mice is in the range of 2 years (see Methods: *Comparison of mouse and human*). The turnover rate of mice organs is substantially higher as suggested by wound healing dynamics. In fact, the model imposes a higher turnover rate (*δ*_*mouse*_ *>* 0.4 per year) such that organ failure can happen at the age of two years (see parameters in Table 1). In the model with age-dependent loss of stem cell division, a 5 weeks rejuvenation treatment induces a substantial effect on organ failure (Figure 6A). In contrast, a 5 weeks treatment of humans had only weak effects even if applied to high turnover organs (Figure 6B). This remains also true when applied to a human organ with even higher and unphysiological turnover rates. This implies, that the positive results found in preclinical trials in mice [3] cannot be expected in human when the same treatment protocol is applied.

**Figure 6:**
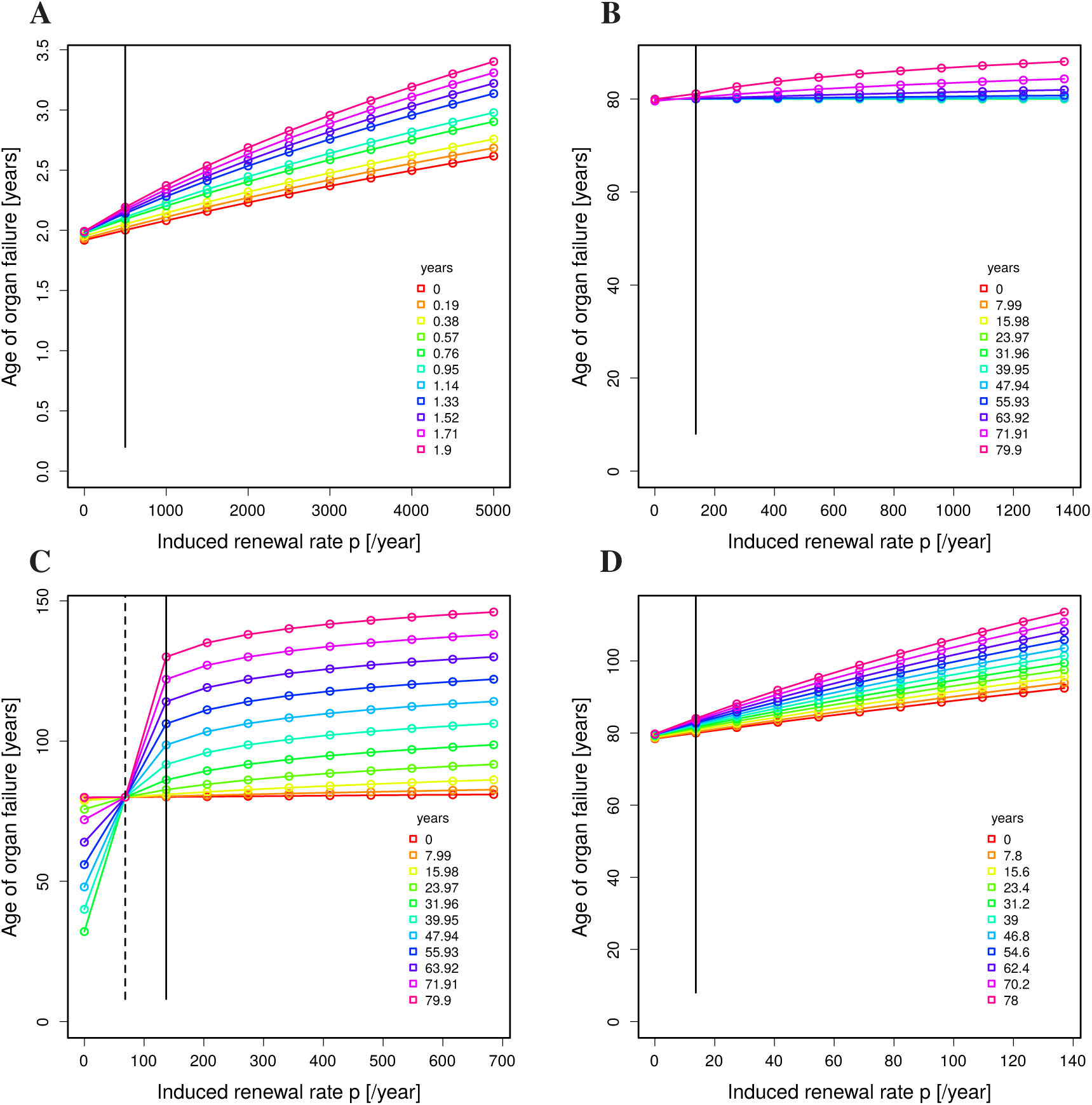
Rejuvenation treatment in mouse and human. (**A**) 5-weeks treatment of mice with life expectancy of 2 years in a model with age-dependent degradation of stem cell replication. (**B**) 5-weeks treatment of humans assuming age-dependent degradation of stem cell replication in a high turnover organ. (**C**) 5-weeks treatment of humans with age-independent degradation of stem cell replication (dashed vertical line) in a high turnover organ. The perfect homeostasis replication rate is induced during treatment (full vertical line). (**D**) 2-years treatment of humans with age-dependent degradation of stem cell replication in a low turnover organ. The parameters are taken from Table 1. The treatment was assumed to induce absolute renewal rates. For relative improvement see Supplementary Figure 5.

The next question was, under which conditions an effect of the treatment is expected to be detectable in humans. Assuming a lifelong constant stem cell replication in high turnover organs, a 5 weeks rejuvenation treatment (inducing *p*_*juvenile*_ in Eq. (11)) leads to a substantial improvement (Figure 6D). This implies that organs with high turnover and with slow ageing like the skin would benefit from short therapies. In contrast, low turnover organs with ageing stem cells like the brain will only exhibit an effect comparable to the one seen in mice when the treatment is prolonged to the scale of years (Figure 6D). Both statement remains true, if one assumes a relative instead of an absolute improvement of stem cell replication (Supplementary Figure 5), however, consistently with the results in Figure 5, the most advantageous age of treatment is altered.

## Discussion

The mathematical analysis of the impact of the transfer of blood from young individuals with the aim to improve stem cell replication predicts that there is an important impact on the delay of organ failure which is payed with a comparably small increase in the risk of developing cancer. The details of improved life time depend on the age of treatment, the turnover rate of the organ, and the mechanism of action of young blood therapy.

The models with age-dependent and age-independent stem cell replication rate have many properties in common. The larger the turnover rate of the considered organ, the more efficiently organ failure is retarded by small improvements in stem cell replication. However, organ failure cannot be further delayed by stronger treatment, i.e. there is natural limit of retardation of organ failure. In organs with higher turnover rates, cancer risk is stable in response to moderate treatments performed at early age. This bears the chance to really delay organ failure without major risk to induce cancer. However, a conservative dosage of the treatment and a subtle analysis of the biological age of the individual are required. At some biological ages, the treatment can induce an unwanted increase of cancer risk.

Further, the treatment strategy has to consider the different turnover rates of the different organs. As the treatment is thought to be provided systemically, all organs are equally affected. But the optimum of treatment dosage and treatment age turned out dependent on the organ turnover rate. As a consequence, the treatment has to be chosen in a rather conservative way, such that none of the organs bears an increased cancer risk, but as many of the organs as possible benefit from delayed organ failure.

The model suggests that the best possible treatment strategy also depends on the mechanistic details of rejuvenation treatment. If the treatment induces an absolute replication rate of stem cells, the age-dependence is similar for all organ turnover rates. However, if the treatment induces an improvement of the replication rate relative to the replication potential of the stem cells at the age of treatment, treatment of older individuals can be without major effect in low turnover organs. Therefore, there is urgent need to clarify the mechanistic details of the effects observed upon rejuvenation therapy [15].

In a mouse parabiosis model, improved tissue homeostasis was found in response to 5-weeks treatments [4, 5]. There is recent evidence that the simple transfer of blood has an impact on tissue homeostasis [8], which opens the possibility to apply a young blood therapy to humans. According to the model results, the treatment schedule identified in mice with positive effect on tissue homeostasis relies on the higher turnover rate of mouse organs. Therefore, a one-to-one application of the treatment schedule to humans will not induce the same degree of rejuvenation, in particular, in low turnover organs like the brain. Longer treatments in the range of years are required, which shall be considered in the interpretation of the outcome of ongoing clinical trials. A negative outcome would not necessarily imply that rejuvenation of human individuals is not possible by blood derived systemic factors. The code developed here may be used to plan clinical trials and to adapt the treatment schedule to the individual state of organs.

The strength of the presented analysis relies on the simplicity of the model based exclusively on generic assumptions and parameters determined by plausible side conditions. The analysis suggests that treatment of ageing by young blood transfer has a realistic chance to increase quality of life in the future. However, the treatment exhibits an age- and organ turnover rate-dependent effect and requires a decent planning.

## Acknowledgement

MMH thanks Georg Pongratz and Rainer Straub for revising the manuscript. MMH was supported by the Helmholtz Initiative on Personalized Medicine (iMED), the German Federal Ministry of Education of Research (eMed: SYSIMIT, SysStomach), and the Human Frontier Science Program (RGP0033/2015).

## Author contribution

MMH developed the research idea and the approach, programmed the code, interpreted the results and wrote the manuscript.

## Methods

### Model equations and analytical solution

The model of tissue homeostasis distinguishes stem cells *S* and tissue cells *T*. Each cell type is classified by the number of accumulated damages with an index *i* for both quantities, i.e. *S*_*i*_ and *T*_*i*_. Model assumptions are described in the main text. The set of ordinary differential equations reads

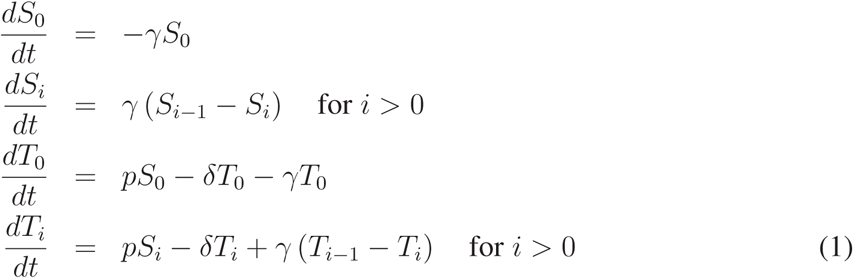

with initial values *S*_*i*_(*t*_0_) = *σ*_*i*_ and *T*_*i*_(*t*_0_) = *τ*_*i*_. *p* is the stem cell replication rate, *γ* is the cell damage rate, and *δ* is the death rate of tissue cells, which corresponds to the turnover rate of the considered organ.

The analytical solution of the model Eq. (1)

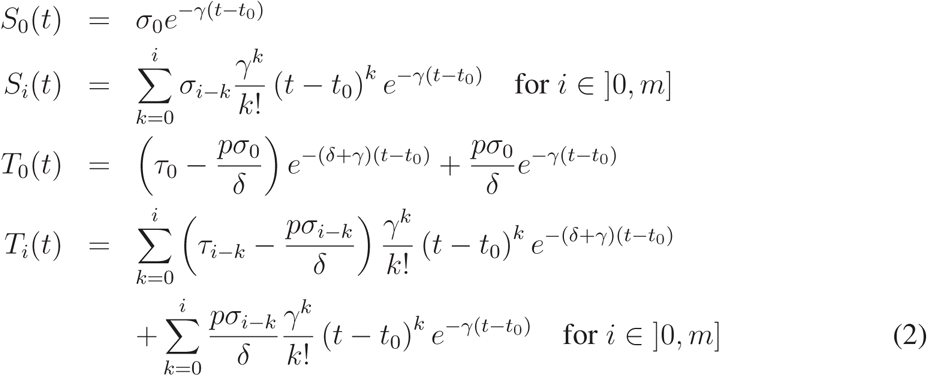

can be proven by complete induction (see Methods: *Proof of the analytical solution*). In the limit of no damage by mutations (i.e. *γ* = 0), the division rate *p* can be determined by the condition of organ failure at the age of Θ = 80 years (see Methods: *Determination of the division rate*). The mutation frequency *γ* is determined by the side condition that the risk of cancer is *ξ* = 30% at the age of Θ = 80 years (see Methods: *Determination of the mutation rate*).

*m* is the maximum number of damages explicitly calculated, with *m* ≫ *c*. Stem and tissue cells with *c* or more accumulated damage are recollected for presentation as

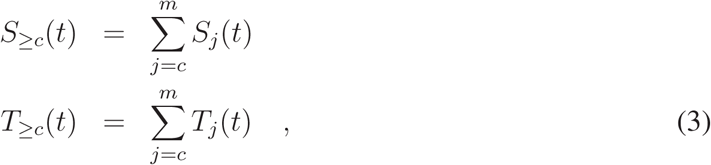

where *m* = 3*c* is chosen sufficiently large such that the results are independent of *m*.

### Determination of the division rate

The division rate *p* can be fixed by the condition that organ failure happens at the age of Θ. Organ failure is assumed to happen when the initial number of tissue cells drops below a fraction *f* = 0.5. It is assumed independent of the number of mutations, thus, Eq. (1) can be solved in the limit of no mutations, i.e. *γ* = 0, which reduces all equations to those indexed with 0. The differential equations for stem and tissue cells read:

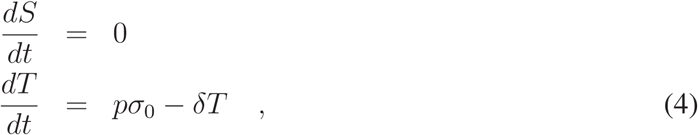

where *S* = *σ*_0_ = *const*. Separation of variables, integration and incorporation of the death condition yields

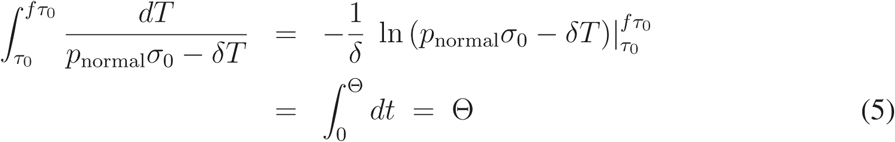

for the unknown division rate *p*_*normal*_. This can be solved for the division rate *p*_*normal*_ resulting in

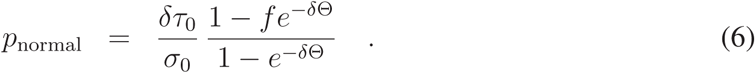

The division rate determined by Eq. (6) is used to set up the dynamics of stem cell division and tissue renewal in normal life without any treatment.

### Determination of the mutation rate

The mutation rate *γ* is determined by the cancer risk *ξ* at the age of life expectancy Θ, where cancer is induced at *c* mutations. Assuming no damage at birth and ignoring more than *c* mutations this condition translates to

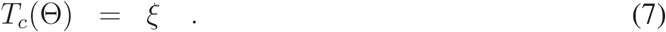

This can be reformulated as

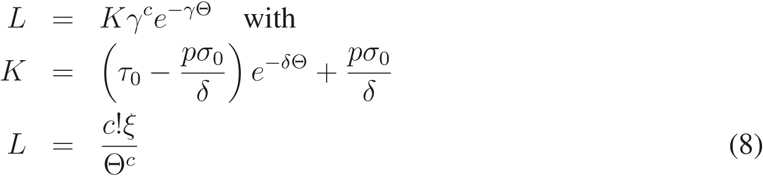

An implicit equation for *γ* of this type is solved by a Lambert W-function as

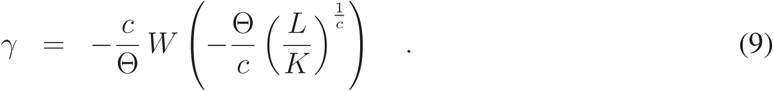

This solution is just an upper bound for *γ* because all contributions of cells with more than *c* mutations were ignored. The condition Eq. (7) with all contributions from cells with *c* or more mutations reads

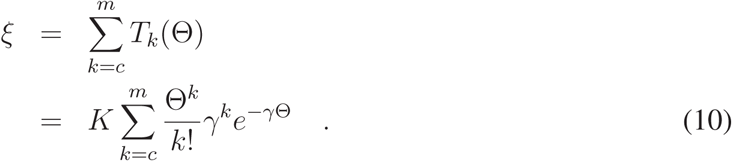

This equation is numerically solved for *γ* where Eq. (9) is used as an initial guess and upper bound.

### Determination of the division rate of perfect homeostasis

Perfect homeostasis implies that any loss of tissue cells by the term *δT*_*i*_ in Eq. (1) is compensated by stem cell division with the term *pS*_*i*_. As this has to happen for all damage levels *i*, it is possible to just consider the sum of all cells, which leads to the condition

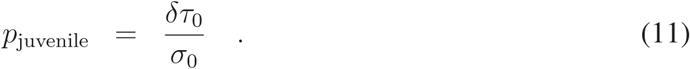

When *p* = *p*_*juvenile*_ for the whole life, organ failure never happens.

### Calculation of the age of cancer risk *ξ*

Given the cancer risk *ξ* and the constant division rate *p*, it is informative to calculate the age Θ_*c*_(*p*), at which the cancer risk reaches *ξ*. This age can be determined from the condition

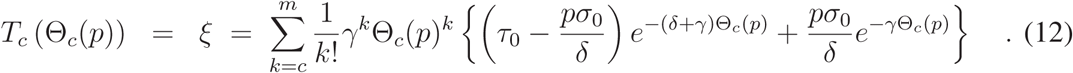

This condition has to be solved for Θ_*c*_(*p*), which was done by numerical approximation of the zero of Eq. (12). The relationship is shown in Figure 2 (blue line).

### Calculation of the age of organ failure

Given the division rate *p* and the initial stem and tissue cell numbers *σ*_0_ and *τ*_0_ it is informative to calculate the age Θ_*f*_(*p*), at which the total tissue cell numbers hit the fraction *f τ*_0_ of organ failure. As the distribution of tissue cells on classes with different levels of accumulated damages does not matter, the condition for organ failure can be calculated in the limit of no damage *γ* = 0. Starting from the analytical solution Eq. (2), the condition reads

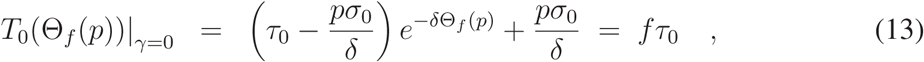

which is solved by

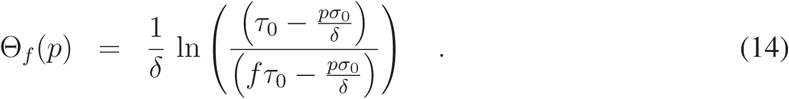

Note that for larger division rates, a physiological solution does not exist in the sense of the argument of the logarithm becoming negative. This relationship is shown in Figure 2 (red line).

### Treatment at particular ages

When the *k*-th treatment with young blood is initiated at a particular age *t*_*k*_ of the individual, the division rate *p* is changed at this particular age. This implies that the cell damage distribution 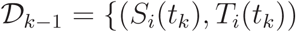 for *i* ∈ [0*, m*]} calculated by the time *t*_*k*_ was used as initial condition for the solution Eq. (2). The development for times *t > t*_*k*_ are then calculated from Eq. (2) with initial condition 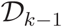, with *t*_0_ = *t*_*k*_, and with the changed *p*. When *p* is changed again, the same procedure has to be repeated. This remains true irrespective of whether the analytical solution Eq. (2) or the numerical solution with age-dependent *p* is used.

### Age-dependent stem cell division rate

Depending on the considered organ, stem cell replication is lost with increasing age. An age-dependent reduction of stem cell replication *p*(*t*) was introduced by replacing each occurrence of *p* in Eq. (1) by *p*(*t*). The solution Eq. (2) is not valid anymore and this system was solved numerically. A Hill-function was used to describe natural loss of stem cell replication as

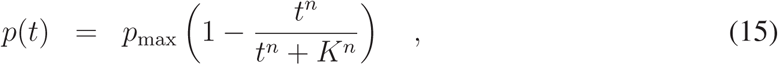

with the replication rate at birth *p*_*max*_ = *p*(*t* = 0) = *p*_*juvenile*_ determined from the assumption of perfect homeostasis Eq. (11). The Hill-coefficient is set to *n* = 2. The age of half replication *K* was chosen such that organ failure occurs at age Θ.

### Comparison of mouse and human

The life expectancy of mice is in the range of two years [16]. With the organ turnover rates *δ* for humans (see Table 1), organ failure at the age of two years cannot be induced in the model (see Eq. (13)). Substantially higher turnover rates with *δ >* 0.4 per year are required to get a solution of the model with positive division rates *p*. At such high turnover rates, organ failure at the age of human life expectancy Θ = 80 years is not achieved (see Eqs. (6) and (13)).

### Robustness of the statements

The sensitivity of the results to changes in the parameter values used in Table 1 and in Eq. (15) is analysed.

The number of mutations per cell associated with induction of cancer *c* was chosen arbitrarily as *c* = 10. For very low *c* the system gets unstable. For larger *c* the behaviour and all described results remain qualitatively unchanged. There is strictly no impact of *c* on the age of organ failure. The impact of treatment on the age of cancer risk *ξ* is further reduced for larger values of *c* (see Supplementary Figure 6). The results based on *c* = 10 can be considered as an upper limit for the increase in cancer risk, as the number of mutations inducing cancer is likely not smaller than 10 but for extreme cases.

Larger *n* in Eq. (15) reduces while smaller *n* increases the overall effect on both, the ages of organ failure and cancer risk *ξ*, in particular, for late treatments (see Supplementary Figure 7). This is consistent with the observation that in the extreme case of constant *p* the effect was larger as well (see Figure 3). A determination of the stem cell activity in human organs over life would allow for a more concrete attribution of therapy effects *in silico* to human organs.

A change in the relative proportion of stem cells *σ*_0_ and tissue cells *τ*_0_ has strictly no influence on the results (see Supplementary Figure 8). The stem cell division rate, by construction, compensates for a different fraction of stem cells. However, the association of the virtual turnover rates with real tissues turnover rates changes.

### Proof of the analytical solution

The ordinary differential equation system Eq. (1) can be solved analytically (see Eq. (2)). The solution can be proven by complete induction, which is sketched here. For the stem cell compartment *S*_*i*_ with *i* ≥ 0 the base case is explicitly calculated. The solutions for *S*_0,1_ are trivial. Let us assume that the general solution Eq. (2) is valid for *S*_*i*_. Then

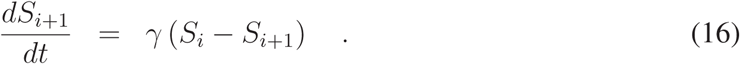

A differential equation of the form

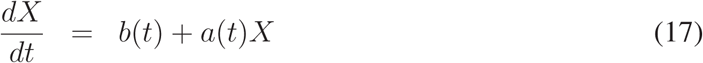

is solved by

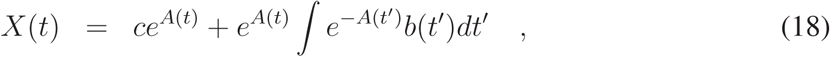

where *A* is a primitive of *a* and *c* is an integration constant. In the case of Eq. (16), we have

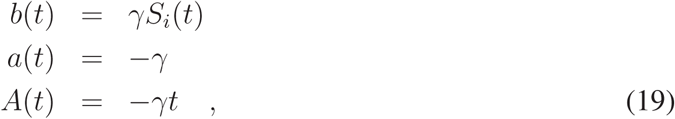

where *S*_*i*_ is taken from the assumed solution Eq. (2). Then

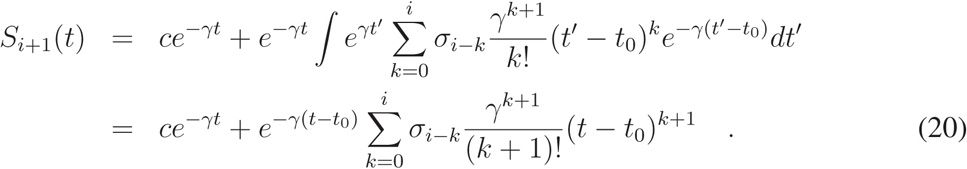

With the initial condition

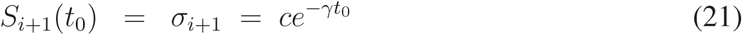

the integration constant is determined as

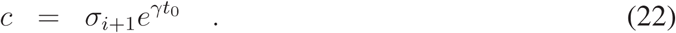

Insertion into Eq. (20) after some algebra yields

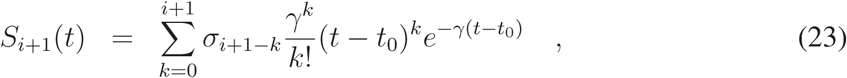

which is what needed to be shown.

For the tissue compartment the base case is calculated for *T*_0_:

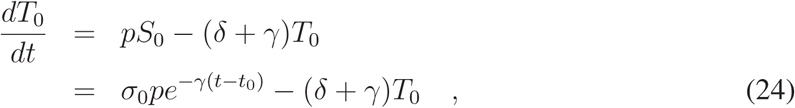

where the explicit solution for *S*_0_ was inserted. This equation is solved with Eq. (18) and the initial condition *T*_0_(*t*_0_) = *τ*_0_ is used to yield

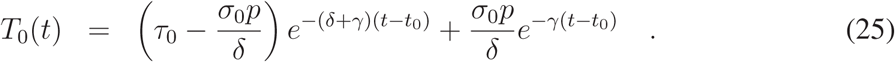

This shows that the explicit solution of *T*_0_ is consistent with the general solution Eq. (2). Now the solution Eq. (2) is assumed valid for *T*_*i*_. Then

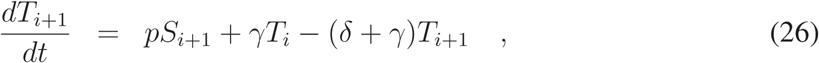

where *S*_*i*+1_ is known from the already proven solution Eq. (2) and *T*_*i*_ can be inserted from the assumed solution. Again Eq. (18) can be applied with

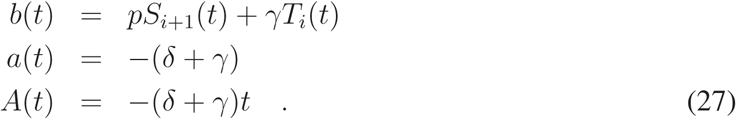

Together with

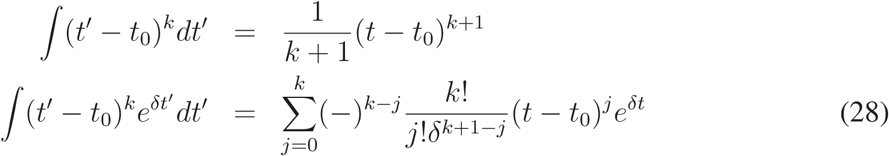

and some algebra, one finds

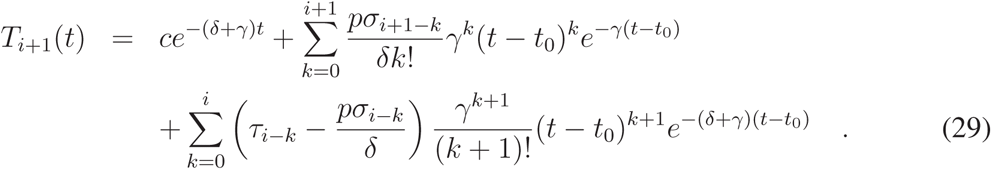

With the initial condition *T*_*i*+1_(*t*_0_) = *τ*_*i*+1_ the integration constant is determined as

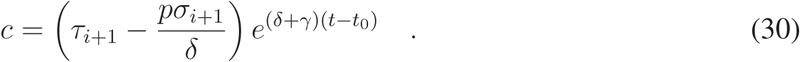

Insertion in Eq. (29), resorting, and renumbering of the sum indices yields:

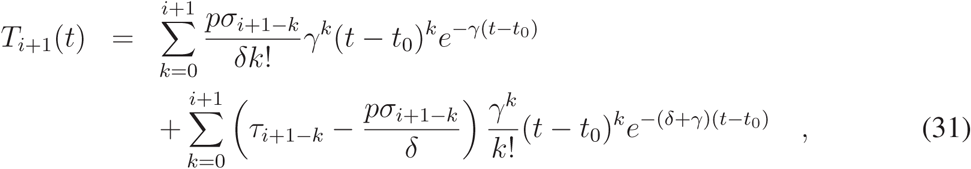

which proves the solution Eq. (2) for all *i*.

## Supplementary figures

**Supplementary Figure 1:**
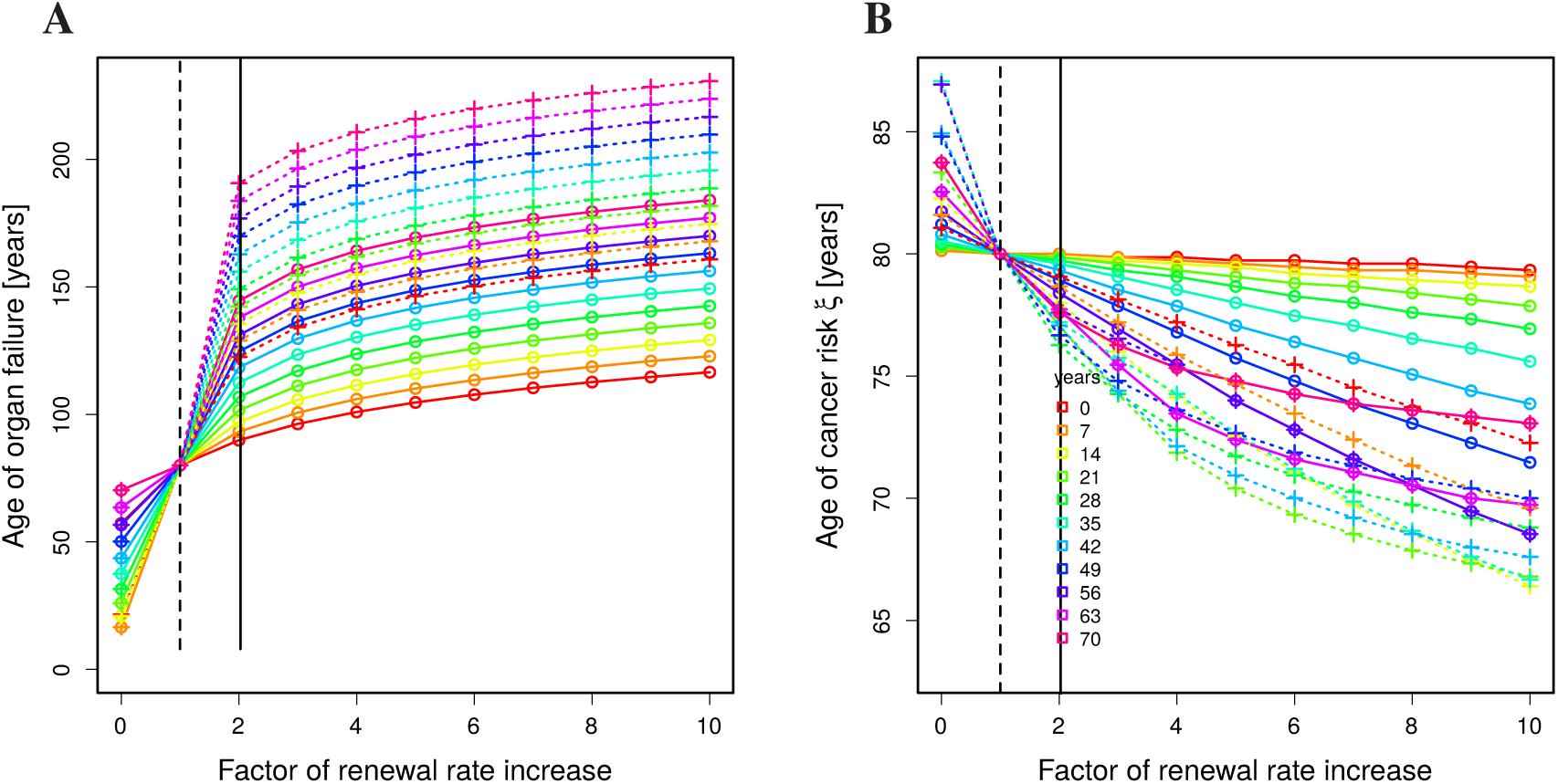
Constant renewal with intermediate turnover rates. Same analysis and representation as in Figure 3 but for organs with intermediate turnover (see Table 1).

**Supplementary Figure 2:**
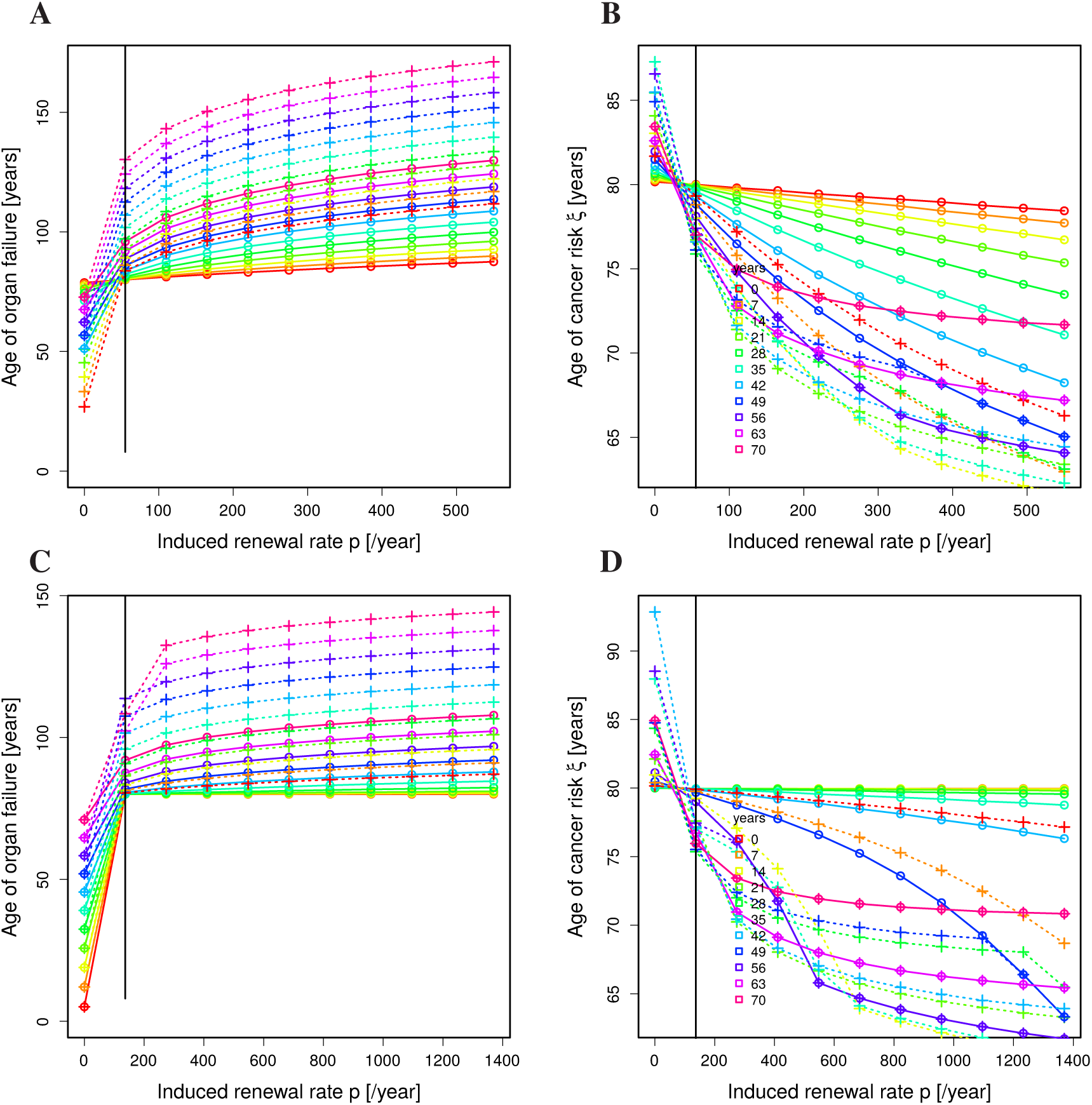
Age-dependent renewal rate for higher turnover organs. Same analysis (i.e. inducing absolute renewal rates by the treatment) and representation as in Figure 5A,B but for organs with intermediate (**A,B**) or high (**C,D**) turnover (see Table 1).

**Supplementary Figure 3:**
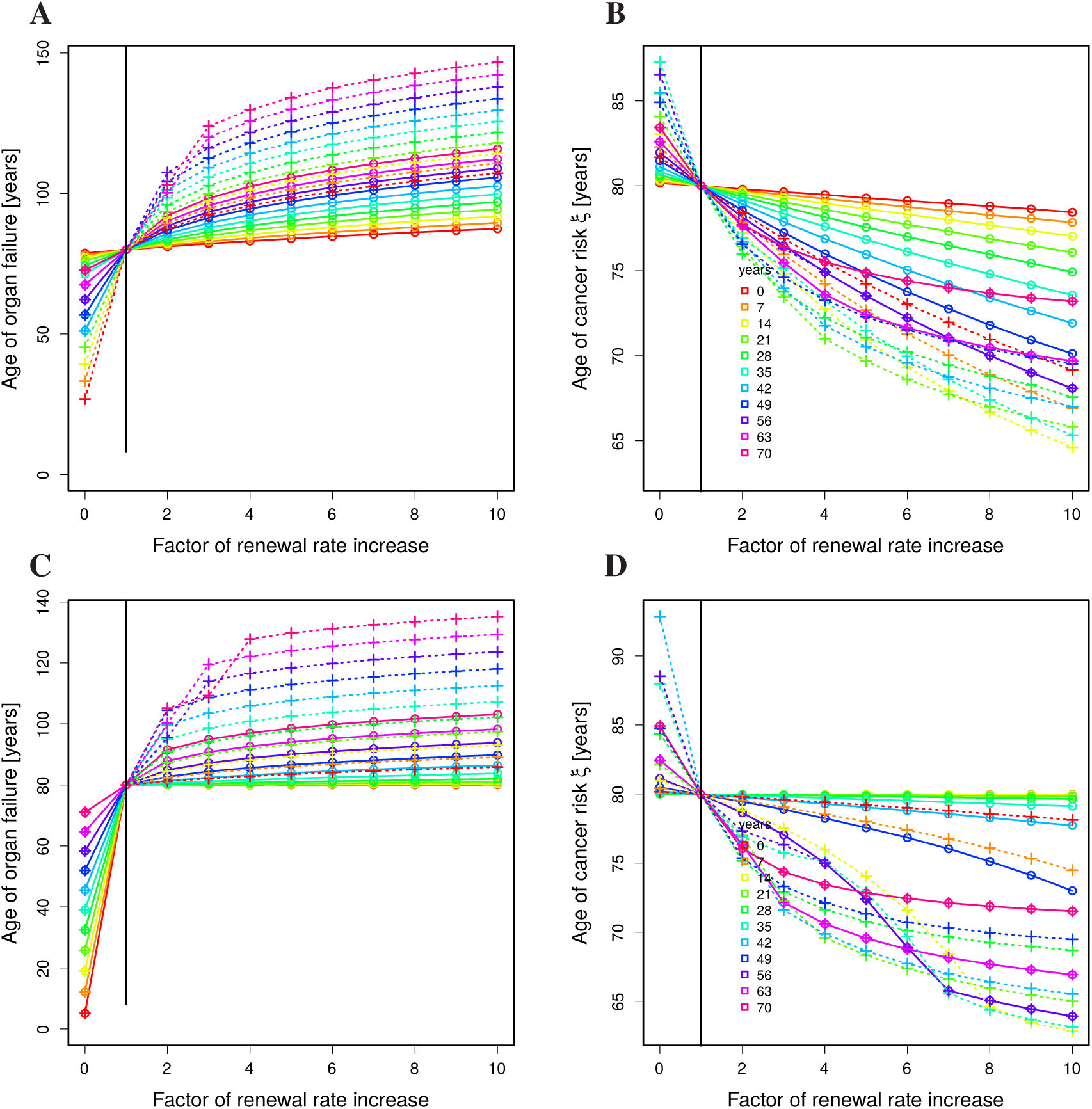
Impact of higher turnover rates. Same analysis (i.e. inducing relative improvements of the renewal rate by the treatment) and representation as in Figure 5C,D but for organs with intermediate (**A,B**) or high (**C,D**) turnover (see Table 1). The hierarchy, that organ failure happens the later the older the treated individual is restored (**A,C**). Cancer risk analysis (**B,D**) is consistent with the case of constant replication rate *p* (compare Figure 3).

**Supplementary Figure 4:**
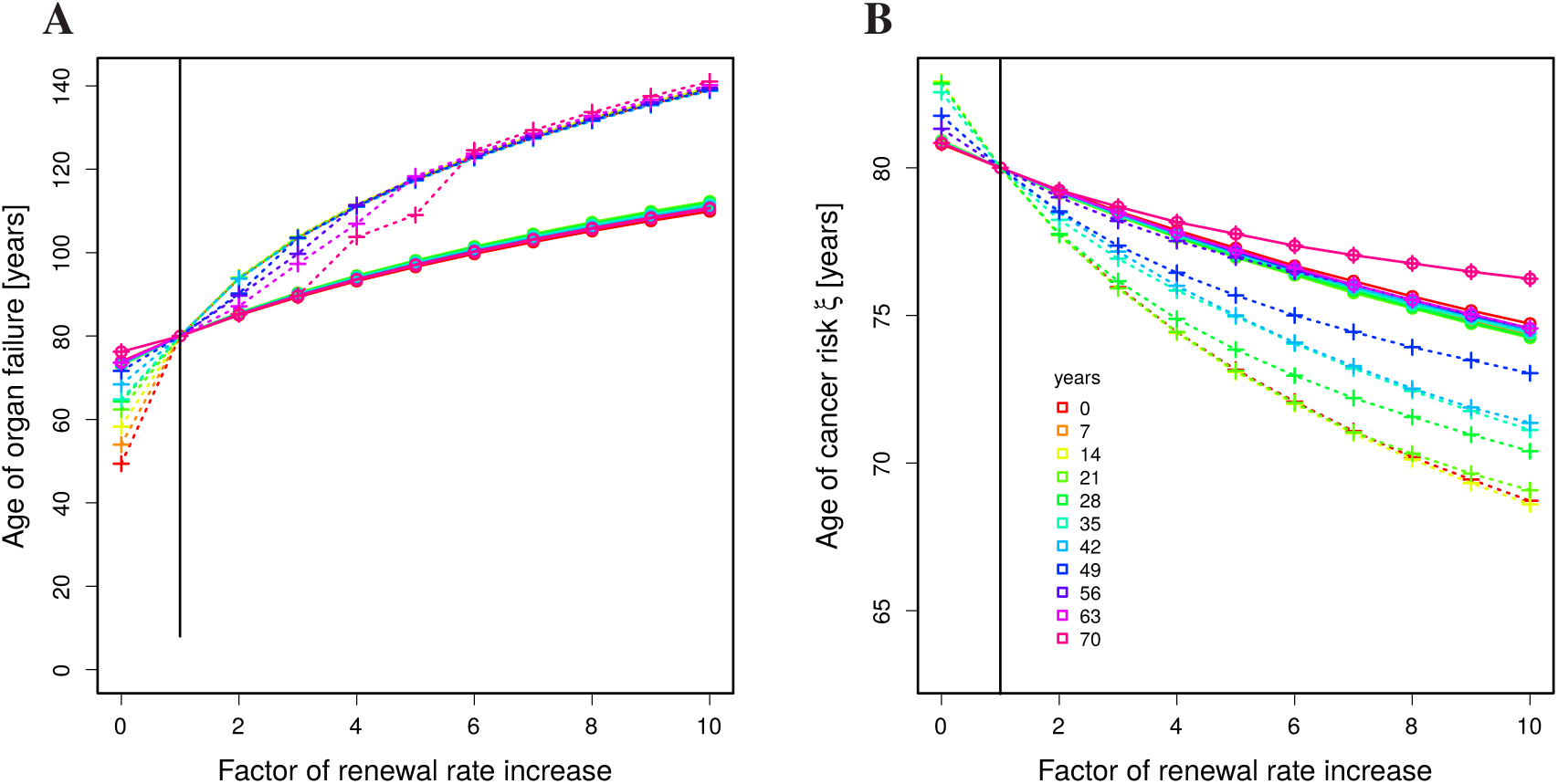
The organ turnover rate associated with age-independent treatment success. Same analysis and representation as in Figure 5 with the critical death rate *δ* = 0.023 per year, at which the age of first treatment becomes irrelevant for the retardation of organ failure. Relative impact of treatment on stem cell replication.

**Supplementary Figure 5:**
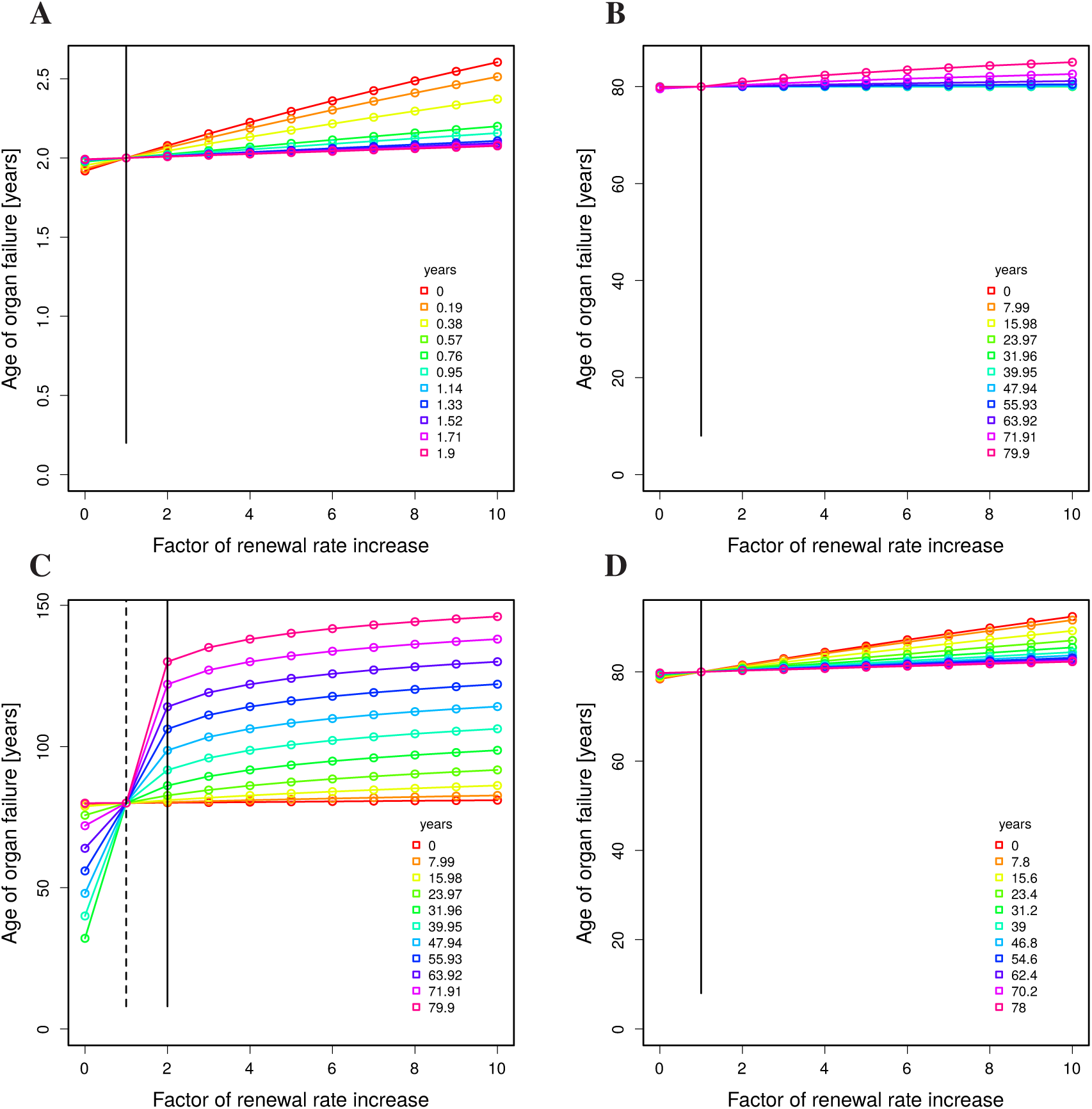
Rejuvenation treatment in mouse and human. Same experiment as in Figure 6 but for relative induced stem cell replication rates (horizontal axes). (**A**) Mouse, age-dependent renewal, treated 5 weeks. (**B**) Human, age-dependent renewal, high turnover organ, treated 5 weeks. (**C**) Human, age-independent renewal, high turnover organ, treated 5 weeks. (**D**) Human, age-dependent renewal, low turnover organ, treated 2 years.

**Supplementary Figure 6:**
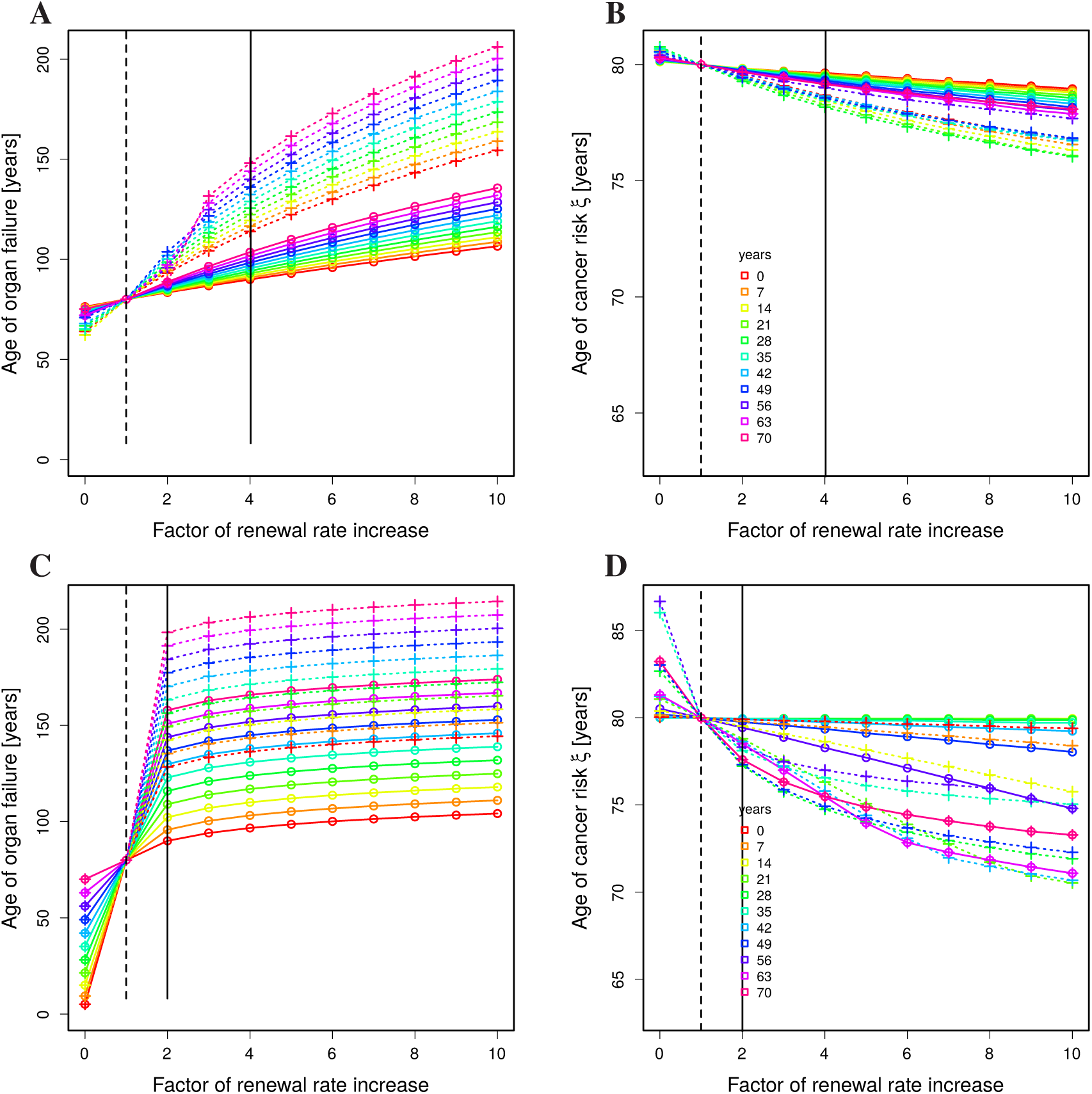
Constant renewal rate with cancer induced at *c* = 20 mutations. Same analysis and representation as in Figure 3 with low (**A,B**) and high (**C,D**) turnover rates (see Table 1).

**Supplementary Figure 7:**
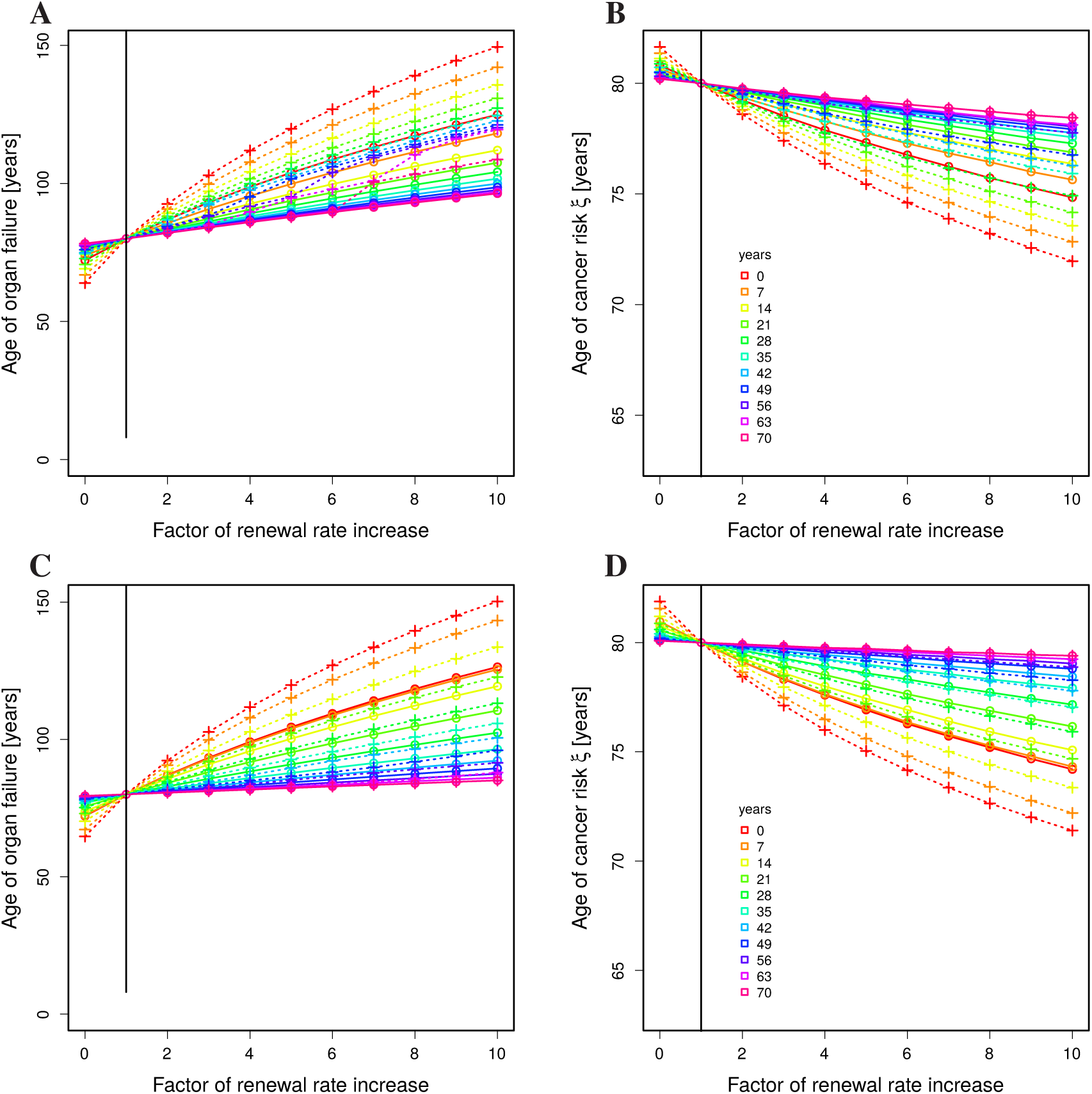
Steep or flat age-dependence of the renewal rate. Same analysis and representation as in Figure 5C,D with flat (*n* = 1.5, **A,B**) and steep (*n* = 3, **C,D**) age-dependence in Eq. (15). Parameters of low turnover rates (see Table 1).

**Supplementary Figure 8:**
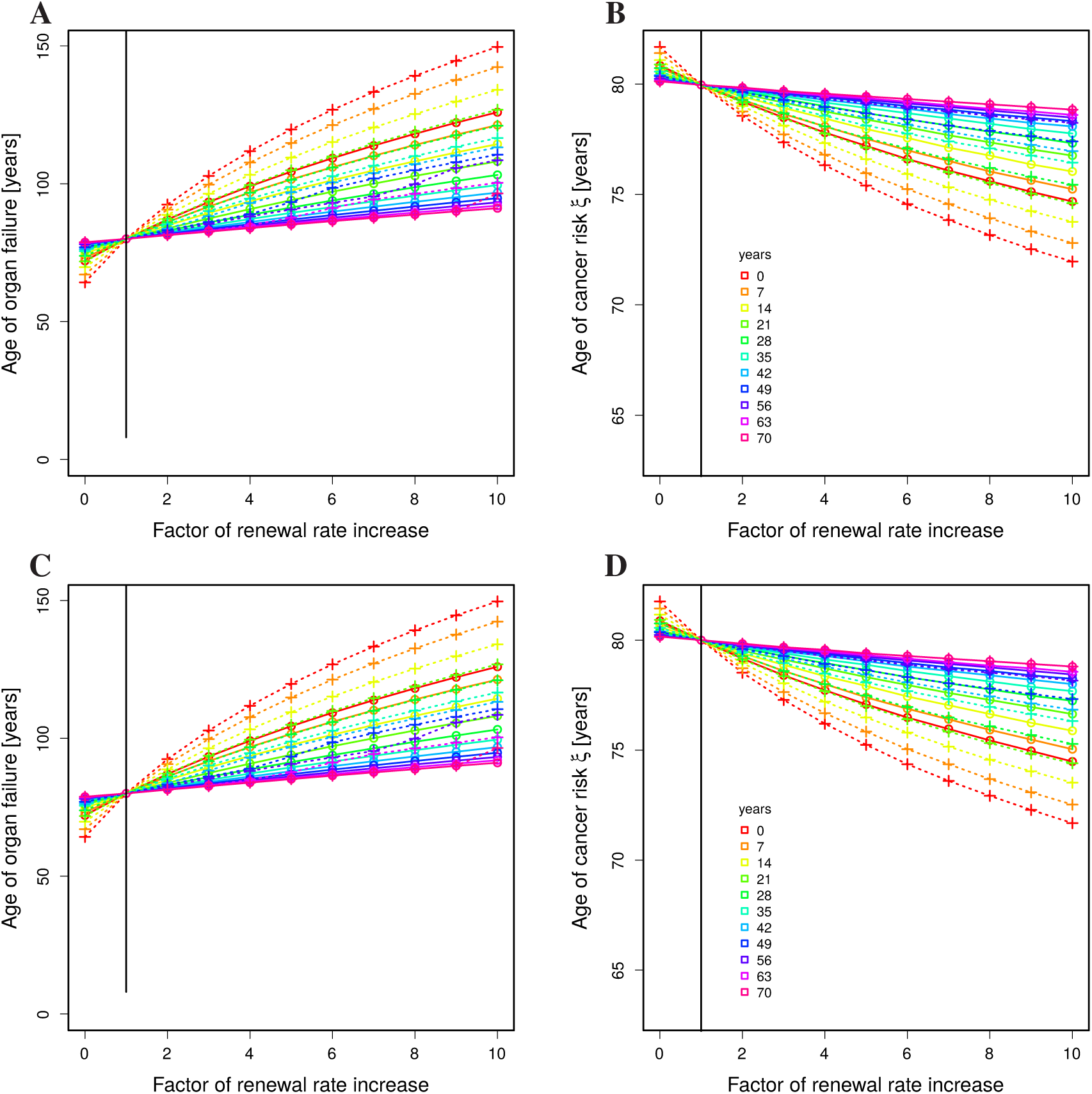
Increased total cell numbers. Same analysis and representation as in Figure 5C,D with 1000 times more cells (**A,B**) and 10 times more stem cells (**C,D**). There is no impact by construction. Using age-dependent renewal rate and relative improvement in 10 years treatments. Parameters of low turnover rates (see Table 1).

## References

1. DeCarolis, N., Kirby, E., Wyss-Coray, T., and Palmer, T. The role of the microenvironmental niche in declining stem-cell functions associated with biological aging. Cold Spring Harb Perspect Med 5, a025874 (2015).

2. Conboy, I., Conboy, M., Wagers, A., Girma, E., Weissman, I., and Rando, T. Rejuvenation of aged progenitor cells by exposure to a young systemic environment. Nature 433, 760–764 (2005).

3. Middeldorp, J., Lehallier, B., Villeda, S., Miedema, S., Evans, E., Czirr, E., Zhang, H., Luo, J., Stan, T., Mosher, K., Masliah, E., and Wyss-Coray, T. Preclinical assessment of young blood plasma for Alzheimer disease. JAMA Neurol 73(11), 1325–1333 (2016).

4. Conboy, M., Conboy, I., and Rando, T. Hereochronic parabiosis: historical perspective and methodological considerations for studies of aging and longevity. Aging Cell 12, 525–530 (2013).

5. Castellano, J., Palner, M., Li, S.-B., Freeman Jr, G., Nguyen, A., Shen, B., Stan, T., Mosher, K., Chin, F., Lecea, L. d., Luo, J., and Wyss-Coray, T. In vivo assessment of behavioral recovery and circulatory exchange in the peritoneal parabiosis model. Sci Rep 6, 29015 (2016).

6. Castellano, J., Kirby, E., and Wyss-Coray, T. Blood-borne revitalization of the aged brain. JAMA Neurol 72(10), 1191–1194 (2015).

7. Wyss-Coray, T. Ageing, neurodegeneration and brain rejuvenation. Nature 539(7628), 180–186 (2016).

8. Rebo, J., Mehdipour, M., Gathwala, R., Causey, K., Liu, Y., Conboy, M., and Conboy, I. A single heterochronic blood exchange reveals rapid inhibition of multiple tissues by old blood. Nat Commun 7, 13363 (2016).

9. Stratton, M., Campbell, P., and Futreal, P. The cancer genome. Nature Reviews 458, 719–724 (2009).

10. Blokzijl, F., Ligt, J. d., Jager, M., Sasselli, V., and Roerink, S. e. a. Tissue-specific mutation accumulation in human adult stem cells during life. Nature 538, 260–264 (2016).

11. Tomasetti, C., Li, L., and Vogelstein, B. Stem cell divisions, somatic mutations, cancer etiology, and cancer prevention. Science 355, 1330–1334 (2017).

12. Giangreco, A., Qin, M., Pintar, J., and Watt, F. Epidermal stem cells are retained in vivo throughout skin aging. Aging Cell 7, 250–259 (2008).

13. Sudo, K., Ema, H., Morita, Y., and Nakauchi, H. Age-associated characteristics of murine hematopoietic stem cells. J Exp Med 192, 1273–1280 (2000).

14. Dykstra, B., Olthof, S., Schreuder, J., Ritsema, M., and Haan, G. d. Clonal analysis reveals multiple functional defects of aged murine hematopoietic stem cells. J Exp Med 208, 2691–2703 (2011).

15. Brunet, A. and Rando, T. Interaction between epigenetic and metabolism in aging stem cells. Curr Opin Cell Biol 45, 1–7 (2017).

16. Graber, T., Kim, J.-H., Grange, R., McLoon, L., and Thompson, L. C57BL/6 life span study: age-related declines in muscle power production and contractile velocity. Age 37, 9773 (2015).

